# A haplotype-resolved chromosomal reference genome for the porcini mushroom *Boletus edulis*

**DOI:** 10.1101/2024.12.09.627560

**Authors:** Etienne Brejon Lamartinière, Keaton Tremble, Bryn T.M. Dentinger, Kanchon K. Dasmahapatra, Joseph I. Hoffman

**Affiliations:** Department of Evolutionary Population Genetics, Faculty of Biology, Bielefeld University, 33501 Bielefeld, Germany; Department of Biology, Duke University, Durham NC, 27707, USA; School of Biological Sciences, University of Utah, Salt Lake City, UT 84112, USA; Natural History Museum of Utah, Salt Lake City, UT 84108, USA; Department of Biology, University of York, Heslington, United Kingdom; Center for Biotechnology (CeBiTec), Faculty of Biology, Bielefeld University, 33615 Bielefeld, Germany; Joint Institute for Individualisation in a Changing Environment (JICE), Bielefeld University and University of Münster, Bielefeld, Germany; British Antarctic Survey, High Cross, Madingley Road, Cambridge CB3 OET, UK

**Keywords:** *Boletus edulis*, ectomycorrhizal fungi (EMF), reference genome assembly, Hi-C, population structure

## Abstract

Haplotype-resolved chromosomal reference genomes are increasingly available for many fungi, offering insights into the evolution of pathogenic and symbiotic lifestyles. However, these resources remain scarce for ectomycorrhizal fungi (EMF), which play crucial roles in forest ecosystems. Here, we used a combination of chromatin conformation capture (Hi-C) and PacBio sequencing to construct a haplotype-resolved chromosomal genome assembly for *Boletus edulis*, an emerging model for EMF research and a prized edible fungus. Our new reference assembly, “BolEdBiel_h2”, derives from a sporocarp sampled in Bielefeld, Germany. The genome assembly spans 41.8 Mb, with a scaffold N50 of 4.1 Mb, and includes 11 chromosome-level scaffolds, achieving near telomere-to-telomere coverage across multiple chromosomes. We annotated 15,406 genes, with a Benchmarking Universal Single-Copy Orthologs (BUSCO) score of 96.2%. Key genomic features such as mating loci, CAZymes, and effector proteins, were mapped across the assembly. As a first application of this new genomic resource, we mapped whole genome resequencing data from 49 genets to investigate the population structure and genetic diversity of the European lineage of *B. edulis*. We identified five distinct genetic clusters and found that high-latitude populations from Iceland and Fennoscandia exhibited greater nucleotide diversity than populations from the United Kingdom and central Europe. Additionally, we discovered a 0.4 Mb inversion on chromosome three and identified several regions of locally elevated nucleotide diversity, which may represent candidates for ecological adaptation. This genomic resource will facilitate a deeper understanding of this ecologically and commercially important wild fungus.

## Introduction

Ectomycorrhizal fungi (EMF) play a crucial role in the functioning of forest ecosystems worldwide. They facilitate nutrient cycling (Read and Perez-Moreno 2003), contribute to carbon sequestration (Anthony et al. 2024), and enhance the growth, immunity and pathogen resistance of nearly 60% of all tree stems on Earth (Steidinger et al. 2019). However, despite their critical role, our understanding of the ecology and evolution of EMF remains limited. This knowledge gap arises partly from the difficulty of studying these predominantly subterranean organisms, which typically cannot be cultured in the laboratory beyond the mycelium stage due to their symbiotic lifestyles.

Over the past three decades, molecular genetic approaches have been pivotal in expanding our understanding of EMF, facilitating both ecological and evolutionary research (Douhan et al. 2011). Classical genetic markers such as RAPDs (Randomly Amplified Polymorphic DNAs), AFLPs (Amplified Fragment Length Polymorphisms) and microsatellite SSRs allowed researchers to distinguish individual genotypes in natural settings, paving the way for studies of clonality and genetic diversity (Bonello et al. 1998; Amend et al. 2009). More recently, advancements in high-throughput sequencing technologies have broadened the scope of EMF research, providing detailed insights into population structure (Branco et al. 2016), gene flow (Tremble, Hoffman, et al. 2023), gene content (Kohler et al. 2015), adaptation (Bazzicalupo et al. 2020) and molecular evolution (Looney et al. 2022).

A key requirement for modern population genomic studies is the availability of high quality, annotated, chromosome-level reference genomes. These resources enable gene discovery (Mariene and Wasmuth 2024) and the characterization of patterns of variation across the genome, including structural variants (Amarasinghe et al. 2020), runs of homozygosity (Brejon Lamartinière et al. 2024) and recombination landscapes (Weissensteiner et al. 2017). New sequencing approaches such as Hi-C (High-throughput Chromosome Conformation Capture;Belton et al. 2012) have been instrumental in improving the contiguity of genome assemblies by allowing sequencing reads to be assembled into phased haplotypes. These techniques have already been used to generate chromosomal reference genomes for several cultured and domesticated fungi (Morin et al. 2012; Engel et al. 2013; Yu et al. 2022; Ma et al. 2023). However, to date, only two chromosomal reference genomes have been published for EMF (Kurokochi et al. 2023; Zhang et al. 2024).

Generating and analyzing long contiguous DNA sequences may represent the next frontier in genomic studies of EMF, as multiple recent studies of cultured and domesticated fungi have uncovered genomic structural variation both among and within dikaryons that may play a role in fungal evolution. For example, (Sperschneider et al. (2023) used phased chromosomal assemblies of the arbuscular mycorrhizal fungus *Rhizophagus irregularis* to demonstrate that separate heterokaryon haplotypes are distinct functional and regulatory units, and can independently modulate the expression host plant genes. Similarly, (Borgognone et al. 2018) showed that DNA methylation in the saprotrophic genus *Pleurotus* can be haplotype-specific and tends to be higher around transposable elements (TEs), where they reduce potentially maladaptive gene expression. As many EMF also have a heterokaryotic life stage, haplotype-resolved chromosomal EMF reference genomes are likely to become important resources for understanding the biology of this important group of fungi.

*Boletus edulis* Bull., known variously as the cèpe de Bordeaux, Steinpilz or porcino, is one of the most charismatic and economically important EMF species worldwide. While the majority of commonly found EMF associate with just one or two host plant genera (Voller et al. 2024), *B. edulis* forms mutualistic associations with diverse plant genera, from the most dominant forest trees of the northern hemisphere ( *Fagus*, *Quercus*, *Pinus*, *Picea*, *Betula*, *Castanea* and *Pseudotsuga*) to alpine miniature shrubs (Treindl and Leuchtmann 2019). It also has a broad geographical distribution spanning Eurasia and North America. For these reasons, *B. edulis* has recently emerged as a promising model system for studying the ecology and evolution of EMF (Hoffman et al. 2020; Tremble et al. 2020; Tremble, Hoffman, et al. 2023; Tremble, Brejon Lamartinière, et al. 2023; Brejon Lamartinière et al. 2024).

The first global-scale population genomic study of *B. edulis* found evidence for seven distinct lineages, which diverged from one another around 1.6–2.7 million years ago (Tremble, Hoffman, et al. 2023). Reference genomes have already been produced for each of these lineages (Tremble, Brejon Lamartinière, et al. 2023), but these vary considerably in contiguity and completeness, and none of them reach the chromosomal level. The most contiguous of these reference genomes, which comprises a total of 38 scaffolds and an N50 of 2.5Mbp 65 scaffolds with an N50 of 2.16 Mbp, is available for the Alaska lineage, while the least contiguous reference genome, which comprises 488 scaffolds with an N50 of 0.17 Mbp, is available for the European lineage. Moreover, these reference genomes were generated using haplotype-unaware methods, which limits the ability to detect structural variants (SVs), especially in highly repetitive genomes (Ebert et al. 2021).

Within the *B. edulis* complex, the European lineage exhibits the widest geographical and ecological distribution, ranging from Mediterranean grasslands to the Scandinavian tundra. This lineage is also associated with the greatest diversity of host species (MyCoPortal, http://www.mycoportal.org/portal/index.php), which is reflected by its expanded symbiosis-related gene repertoire (Tremble, Brejon Lamartinière, et al. 2023). Developing a chromosomal reference genome for this lineage would allow for a more in-depth exploration of population structure, local adaptation, and host specialization, not only across Eurasia, but also on a broader scale. This is particularly important in the context of climate change, as many forests of the northern hemisphere are being strongly impacted by warmer and drier conditions (Gazol et al. 2017). A deeper understanding of forest adaptation is essential, and it must include EMF, which play a vital role in helping trees mitigate climate-induced stress.

In this study, we combined highly accurate long-read Pacbio HiFi sequencing with Hi-C to generate a haplotype-resolved chromosomal genome assembly for the European *B. edulis* lineage. We predicted key genomic features including mating loci, carbohydrate-active enzymes (CAZymes), transposable elements (TEs) and overall gene content. Additionally, we investigated structural and gene copy variation within the reference individual and explored longer-term patterns of synteny with *Suillus bovinus (*Zhang *et al. 2024)*. To further contextualize the reference genome, we mapped short-read data from 49 European samples to examine patterns of population genetic structure within the European *B. edulis* lineage.

## Materials and methods

### Sporocarp tissue sampling

A total of 15g of tissue from the inner cap flesh of a *B. edulis* sporocarp was collected from a *Fagus* woodland in Bielefeld, Germany. Tissue from this sample was archived in the the Senckenberg Museum of Natural History, Görlitz, under the reference GLM-F139661. It was sequentially frozen at 4°C for 30 min, then at -20°C for 30 min, before being stored at -80°C. DNA isolation, library preparation, sequencing and genome component prediction were performed by BMKgene as described below.

### Genomic DNA isolation and sequencing

DNA was extracted from sporocarp tissue using a QIAGEN® Genomic-tip 20G kit. To generate a highly-contiguous assembly, long-read sequencing was conducted as follows. Libraries were constructed according to PacBio standard protocol and sequenced on a PacBio Sequel II platform. The resulting raw circular consensus sequences (CCS) were quality-filtered using smrtlink v12 with the parameters --min-passes 5 –min-rq 0.9, assembled into scaffolds using Hifiasm v0.12 (Cheng et al. 2021) and corrected using Pilon v1.17 (Walker et al. 2014).

Hi-C libraries were constructed as follows. Cross-linking was performed on isolated DNA to preserve the three-dimensional structure of the chromatin and to retain chromatin interactions. The cross-linked DNA was digested with Dpn II and the resulting fragments were repaired with biotin-labelled nucleotides. DNA interactions were stabilized by ligation to form circular structures. The cross-linking was then reversed and the biotin-labeled fragments were captured using streptavidin beads. The purified fragments were inspected using an Agilent 2100 fragment analyzer (Nachamkin et al. 2001) and their concentrations were determined using q-PCR. The libraries were sequenced on an Illumina NovaSeq using paired-end 150 bp reads. The resulting paired-end Hi-C reads were filtered using HiC-Pro v2.10.0 (Servant et al. 2015), separated for each haplotype by integration to the long reads using Hifiasm v0.12 and aligned to the long-read assembly using bwa v0.7.10 (Li and Durbin 2009). The contigs were then clustered, ordered and oriented using Lachesis (Burton et al. 2013). To visualize chromatin interactions and genome contiguity, we generated a chromatin contact heat-map using the command-line version of Juicer v1.6 (Durand et al. 2016)

### Genome annotation

A repeat database was constructed using a combination of LTR_FINDER v1.05 (Xu and Wang 2007), MITE-Hunter (Han and Wessler 2010), RepeatScout v1.0.5 (Price et al. 2005) and PILER-DF v2.4 (Edgar and Myers 2005). The database was sorted with PASTEClassifier (Hoede et al. 2014) and merged with the Repbase database (Bao et al. 2015). RepeatMasker v4.0.6 (Smit et al, 2015) was then used to predict repeat elements in the fungal genome based on this combined database. Gene prediction was performed through both *de novo* and homology-based approaches. The *de novo* prediction utilised Genscan (Burge and Karlin 1997), Augustus v2.4 (Stanke et al. 2004), GlimmerHMM v3.0.4 (Majoros et al. 2004), GeneID v1.4 (Alioto et al. 2018) and SNAP (version 2006-07-28) (Johnson et al. 2008). For homology-based prediction, GeMoMa v1.3.1 (Keilwagen et al. 2019) was used to identify homologous protein coding genes. The results from both approaches were integrated using EvidenceModeler v1.1.1 (Haas et al. 2008). Non-coding RNA was predicted using tRNAscan-SE -version 2.0 (Chan and Lowe 2019) for tRNA, and Infernal 1.1 (Nawrocki and Eddy 2013) for other non-coding RNAs. Pseudogenes were identified by aligning homologous genes from the predicted protein list and the Swiss-Prot database (Bairoch and Apweiler 2000) to the genome using GenBlastA v1.0.4 (She et al. 2009), followed by the detection of early termination and frameshift mutations with GeneWise (Birney et al. 2004). Gene clusters were identified using antiSMASH v6.0.0 (Medema et al. 2011). Carbohydrate-active enzymes (CAZymes) were annotated using hmmer -version (Finn et al. 2011) based on the CAZy database (Cantarel et al. 2009). Effector proteins were predicted using EffectorP v3.0 (Sperschneider et al. 2016).

### Haplotype comparisons and evolutionary context

To investigate the quality of the alternative haplotypes, we identified the positions of conserved Benchmarking Universal Single-Copy Orthologs (BUSCO) genes from the basidiomycota_obd10 database using BUSCO v5.7.1 (Simão et al. 2015) separately for each haplotype. We also ran gene and TE prediction on each haplotype using Funnanotate v1.8.17 (Palmer and Stajich 2023) a and EDTA v2.2.0 (Ou et al. 2019), respectively. To evaluate mapping coverage, we aligned sequencing reads from 49 European *B. edulis* samples to both haplotypes using the mem2 algorithm from bwa (https://github.com/bwa-mem2/bwa-mem2) with default parameters. These sequencing data included previously published Illumina MiSeq, HiSeq and NovaSeq data from 45 samples (Tremble, Brejon Lamartinière, et al. 2023) as well as four additional samples from Bielefeld, Germany, that were 150bp PE sequenced to 30x coverage on a BGI T7 platform. To compute the average depth of coverage per position, we used a 500 kbp sliding window with a 50 kbp step, combining the VCFtools command –site-mean-depth (Danecek et al. 2011) and the GenomicRanges Grange function in R (Lawrence et al. 2013).

After characterizing and comparing the two haplotypes, as described in the Results below, we selected haplotype two for all subsequent analyses. We localized the mating loci by blasting the MATa locus identified by (Tremble, Brejon Lamartinière, et al. 2023) and the annotated STE3 genes from the MATb locus in the *B. edulis* bed1 genome accessed from JGI Mycocosm (Miyauchi et al. 2020) against our reference genome using BLAST 2.2.31 (Altschul et al. 1990). We also checked for the presence of telomeric sequences and located them in the genome using the Telomere Identification Toolkit TIDK (Brown et al. 2023). Finally, we conducted a synteny analysis between the BUSCO genes from the basidiomycota_obd10 database predicted in our reference genome and those from the chromosomal assembly of *S. bovinus* (Zhang et al. 2024) as described above.

### European population structure and genetic diversity

As a first application of this newly generated resource, we analyzed patterns of population genetic structure and genome-wide diversity within the European *B. edulis* lineage. After mapping short-read data from 49 genets to haplotype two of the reference genome following the workflow described above, variants were called using a three-step approach implemented in GATK (McKenna et al. 2010). First, variants were called with HaplotypeCaller, then the files were aggregated into a single database using GenomicsDBImport, and finally, an all-site unfiltered multi-sample VCF file was generated with GenotypeGVCF. This file was then filtered for single nucleotide polymorphisms (SNPs) with a minor allele frequency ≥ 5%, mapping quality ≥ 15, missing genotypes ≤ 20%, a minimum depth of coverage of five and a maximum of 100 using VCFtools. We then computed a principal component analysis (PCA) of the data using Pcadapt (Luu et al. 2017) on the filtered VCF. In addition, we calculated nucleotide diversity (π) in windows of 10 kb with a step size of 1 kb across the genome. This analysis was conducted separately for individuals from different geographical regions with a sample size greater than five, using the VCFtools –window-pi and –window-pi-step commands. The all-site VCF included invariant loci, which were pruned as described previously, while only excluding the minor allele frequency filter.

## Results and Discussion

### Genome assembly quality

The PacBio-generated scaffolds achieved a 100% Hi-C anchoring rate for both haplotypes, resulting in two chromosome-level assemblies consisting of 11 chromosomes each (“BolEdBiel_h1” and “BolEdBiel_h2” respectively, Figure 1a, Figure S1, see Table 1 and Table S1 for details). The high quality of these assemblies is indicated by an excellent chromosome contiguity and their elevated BUSCO scores, which exceeded 96.6% for both haplotypes, with a slightly higher score for haplotype two. Genome sizes for the two haplotypes were 49 Mbp and 41 Mbp respectively, with both exhibiting a GC content of 49.7%. However, the scaffold N50 was slightly lower for haplotype one (3.8 Mbp compared to 4.1 Mbp for haplotype two). This discrepancy is partly due to the fact that over 15% of haplotype one’s total length comprised unanchored scaffolds. When mapping short read sequencing data from 49 European individuals to this haplotype, coverage for all but one unplaced scaffolds was very low (mean = 2.1 versus 24.9 respectively, Figure 1b), reflecting their high repetitive content.

**Figure 1.**
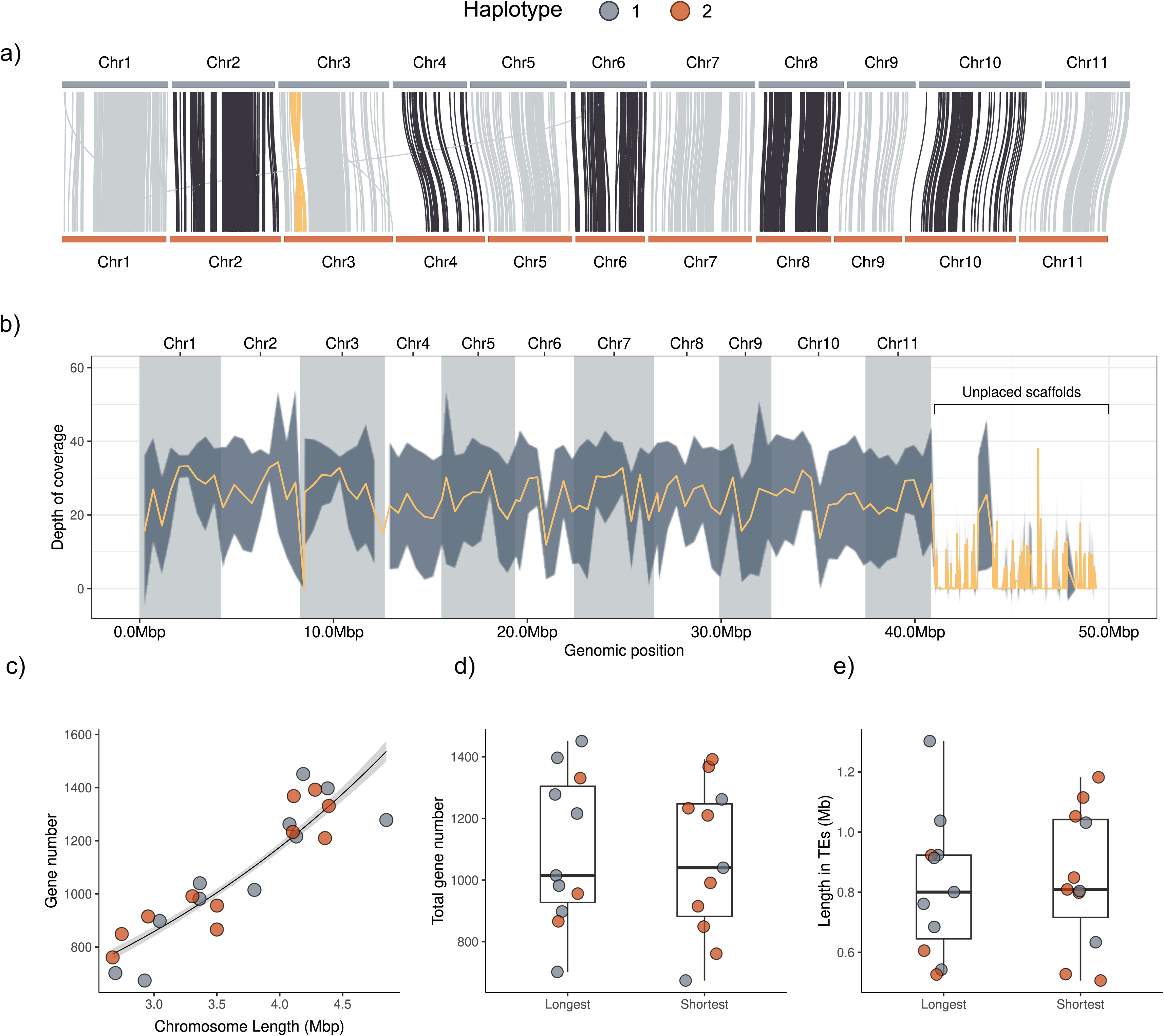
Comparison of the two *B. edulis* haplotype assemblies. a) Synteny between the two haplotypes of each pair of homologous chromosomes. To facilitate visualization, we reversed the alignments of chromosomes five, six, seven and nine from haplotype one, and highlighted the inversion on chromosome three in yellow. b) Depth of mapping coverage of whole-genome resequencing data from 49 genets from the European *B. edulis* linage to haplotype one. The orange line represents the mean mapping coverage, with the dark grey shaded region representing the 95% confidence interval. The chromosomes are represented by alternating grey and white blocks. c) The relationship between chromosome length and gene number. The black line represents the fit of a poisson regression with the grey shaded region representing standard error. Panels d) and e) show comparisons of gene number counts and TE content between the longest and shortest haplotypes respectively. In panels a), c), d) and e), haplotypes one and two are colour coded in grey and orange respectively.

**Table 1:** Quality metrics for the two *B. edulis* haplotype assemblies. BUSCO scores are given for: Complete (C), Complete and single-copy (S), Complete and duplicated (D), Fragmented (F), Missing (M).

| Haplotype | Length (bp) | Chromosomes | Scaffolds | Contigs | Contig N50 (bp) | Scaffold N50 (bp) | n BUSCOs (%) | %GC content |
| --- | --- | --- | --- | --- | --- | --- | --- | --- |
| 1 | 49,135,892 | 11 | 311 | 321 | 3,363,843 | 3,798,680 | C:1704 (96.6%)<br>S: 1630 (92.4%)<br>D: 74 (4.2%)<br>F: 19 (1.1%)<br>M: 41 (2.3%) | 49.7 |
| 2 | 41,815,337 | 11 | 73 | 90 | 2,952,305 | 4,104,162 | C:1709 (96.8%)<br>S: 1643 (93.1%)<br>D: 66 (3.7%)<br>F: 17 (1%)<br>M: 38 (2.2%) | 49.7 |

We found a strong positive association between chromosome length and gene number (z = 30.39, *p* < 0.001, Figure 1c). However, this relationship did not hold when comparing homologous chromosomes. In most cases, haplotype one was longer than haplotype two (Figure 1a,d), yet there were no significant differences between homologous chromosomes in terms of gene count (Figure 1d) and the abundance of TEs (Figure 1e). Based on these results and the slightly better quality metrics for haplotype two, including higher scaffold N50 and BUSCO scores, along with the low coverage of haplotype one-specific contigs, we selected BolEdBiel_h2 as the preferred reference genome for *B. edulis*. Nonetheless, both haplotypes have excellent quality metrics and, given there is no linkage across chromosomes, the choice of haplotype is somewhat arbitrary. Therefore, both assemblies could potentially be used for research in this species, particularly for pan-genomic or structural variation analyses.

The synteny analysis also uncovered a 0.4 Mbp inversion on chromosome 3 (Figure 1a). We confirmed this inversion by assessing PacBio sequence mappings rates using the integrative genomics viewer (Thorvaldsdóttir et al. 2012). Inversions can play important roles in evolution by limiting or suppressing local recombination, which helps to maintain genetic diversity and adaptative potential, and can also drive population divergence and speciation (Hoffmann and Rieseberg 2008). In extreme cases, inversions have been found to induce shifts from mutualism to pathogenesis (Somvanshi et al. 2012). Consequently, further research into this structural variant might help to illuminate on the genetic mechanisms underlying the host adaptability and ecological flexibility of *B. edulis*.

### Genome content

Focusing on BolEdBiel_h2, we predicted a total of 15,406 genes with an average gene length of 1.7kb, which together account for approximately 53.4% of the total genome length (41.8Mbp). Additionally, we identified 11,890 repetitive sequences, representing 35.4% of the genome. Telomeric repeats, (TTAGG)n, were identified in eight of the 11 chromosomes. Telomere-to-telomere assemblies were achieved for chromosomes five and ten (Figure 2), which is a rarity in fungi. Telomeres have been linked to several important biological processes. For example, in the fungal pathogen *Pyricularia*, telomere-adjacent regions are enriched in genes involved in host adaptation genes (Rahnama et al. 2021). Similar patterns could be expected in EMF, potentially offering new insights into the mechanisms underlying host adaptation.

**Figure 2.**
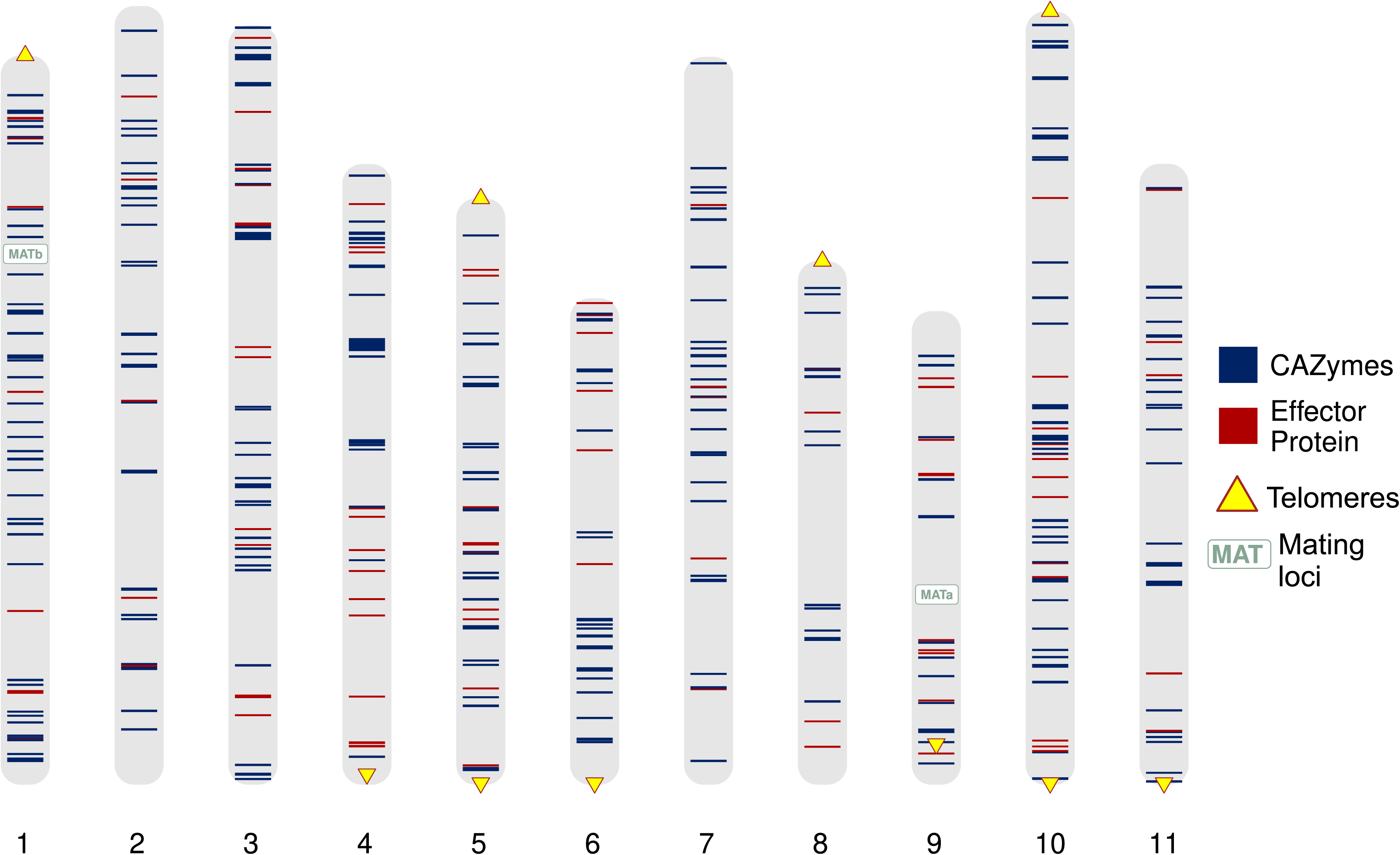
The distribution of key features in the *B. edulis* reference genome. Shown are the genomic locations (in haplotype two) of mating loci, CAZymes, effector proteins and telomeres.

We sought to identify the genomic locations of carbohydrate-active enzymes (CAZymes), which have evolved in fungi to degrade carbohydrates such as lignin and cellulose. While typically associated with saprotrophism, CAZymes are also present in reduced numbers in EMF where they may support nutrient acquisition (Gong et al. 2023). We annotated 413 genes as CAZymes, representing 2.7% of the total gene content. This is comparable to the 2.2% reported for *S. bovinus*, another EMF species (Zhang et al. 2024). These genes were unevenly distributed across chromosomes, with chromosome nine carrying only 15 CAZymes compared to 52 on chromosome one (Figure 2). We also identified genomic regions where multiple CAZymes clustered together. For example, on chromosome four at position 14,673,340, four CAZymes were found within a 54 kbp region (Figure 2). These clusters may represent remnants of an ancestral pre-symbiotic lifestyle, as similar gene arrangements have been observed in wood-decaying fungi (Li et al. 2017). Further exploration of these CAZyme clusters could therefore provide insights into how EMF adapt to diverse environmental conditions.

Effector proteins are secreted by pathogenic fungi to manipulate host defenses during colonization (Li et al. 2024). They have also been shown to play a similar role during the early establishment of mycorrhizal symbioses, facilitating fungal colonization of host roots (Li et al. 2024). However, this process remains poorly understood, and only a small number of EMF effectors have been linked to plant gene expression pathways (Plett et al. 2011; Daguerre et al. 2020). Identifying these genes in the reference genome is an important first step towards understanding their function and regulation in *B. edulis*. We identified a total of 116 effector proteins. Consistent with observations in pathogenic fungi, where effector genes are often clustered in regions of rapid evolution, such as near telomeres (Zaccaron et al. 2022), we found that effector proteins were frequently located near one another and in proximity to telomeric regions. Specifically, we observed this pattern on chromosomes four, five, nine, and ten (Figure 2).

Mating type (MAT) loci play a crucial role in determining sexual compatibility and reproduction in fungi (Coelho et al. 2017). In this tetrapolar species, we identified both mating loci: MATa on chromosome nine and MATb on chromosome one (Figure 2). Tetrapolar species are generally considered less prone to selfing than bipolar species. In line with this, a recent genomic study of *B. edulis* found no evidence of recent inbreeding (Brejon Lamartinière et al. 2024), despite the presence of locally elevated relatedness (Hoffman et al. 2020). This suggests that the mating loci may contribute towards the maintenance of high levels of heterozygosity and genetic diversity within *B. edulis* populations. Future studies could use pedigrees to model the involvement of MAT loci in mating outcomes and inbreeding / outbreeding dynamics.

### Patterns of synteny

To provide phylogenetic and evolutionary context, we investigated the patterns of synteny between *B. edulis* and *S. bovinus* based on shared BUSCO genes. While both species belong to the order Boletales, *S. bovinus* is part of the family Suillaceae, which diverged from Boletaceae approximately 115 million years ago (Wu et al. 2022). Despite sharing an ectomycorrhizal lifestyle with *B. edulis*, *S. bovinus* has a more restricted host range, primarily associating with Pinaceae (Zhang et al. 2024). Surprisingly, we observed high levels of synteny between these species (Figure 3), despite their considerable phylogenetic distance, differences in chromosome numbers, and the structural and size variation present between homologous *B. edulis* haplotypes (Table S1). Notably, large BUSCO gene segments on chromosomes one and eight of *B. edulis* showed structural conservation with chromosomes three and six of *S. bovinus* (Figure 3). Generating additional chromosomal reference genomes for other EMF species would provide deeper insights into genome evolution through comparative chromosomal synteny analyses across multiple species.

**Figure 3.**
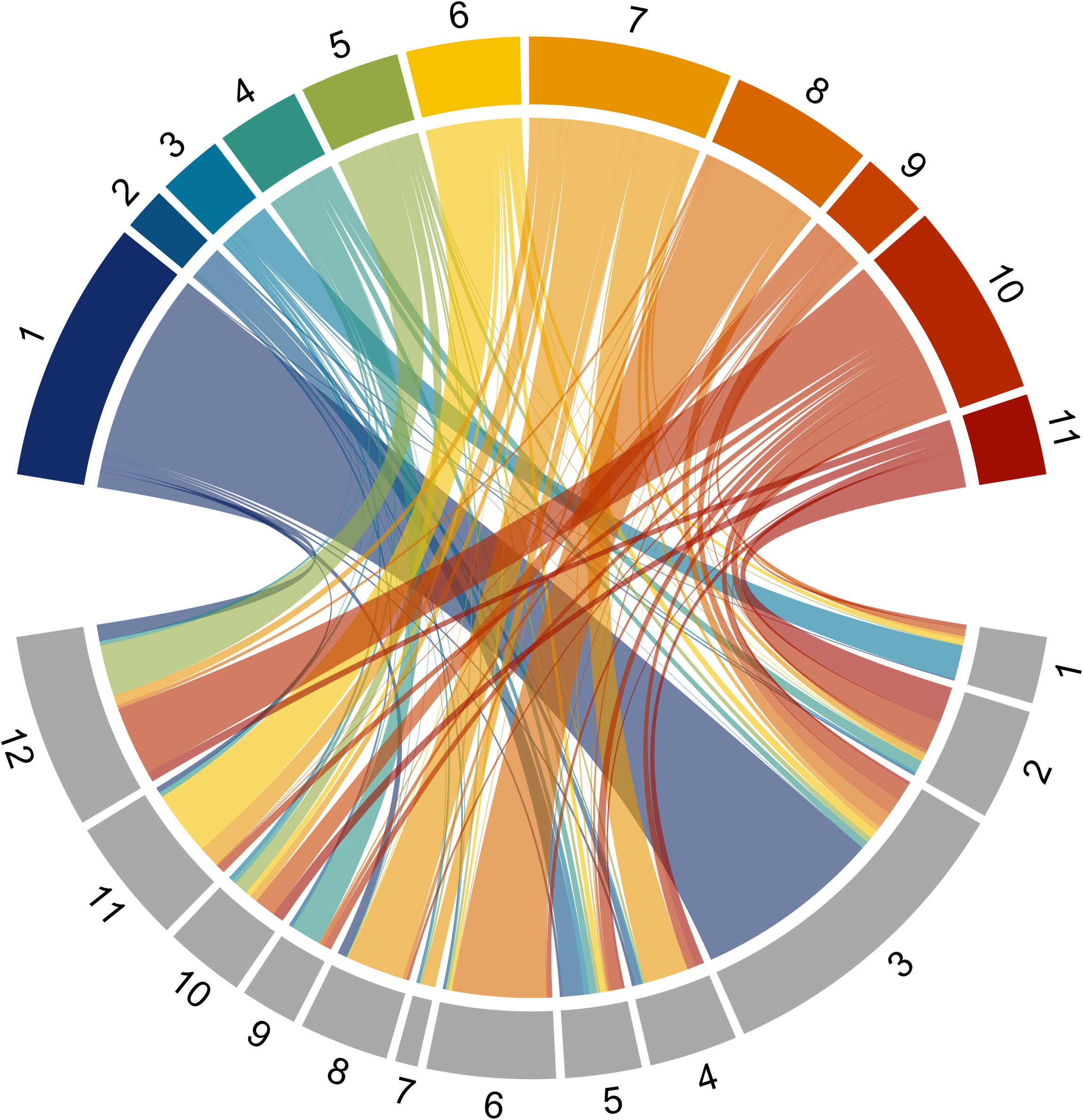
Patterns of chromosomal synteny between *B. edulis* and *S. bovinus*. The circos plot depicts alignments of BUSCO genes from haplotype two of *B. edulis* (top) to the reference genome of *S. bovinus* (below). The *B. edulis* chromosomes are shown in colour and the *S. bovinus* chromosomes are shown in grey.

### Population structure and genetic diversity

We analyzed short-read data from a total of 49 genets (Figure 4a, Tremble, Hoffman, et al. 2023; Brejon Lamartinière et al. 2024, and this manuscript) to genotype 915 929 SNPs for the assessment of population structure and genetic diversity within the European *B. edulis* lineage. In the principal component analysis (PCA), some clustering by geography was observed, corresponding to United Kingdom, Fennoscandia, Iceland, Central Europe, and Southern Europe / Russia respectively (Figure 4b). Notably, a single genet sampled in North America, which was assigned to the European lineage by Tremble, Hoffman, et al. (2023), clustered in the PCA alongside the Fennoscandian and Central European samples (Figure 4b). We speculate that this individual might have been introduced to America from a tree plantation originating in northern Europe.

**Figure 4.**
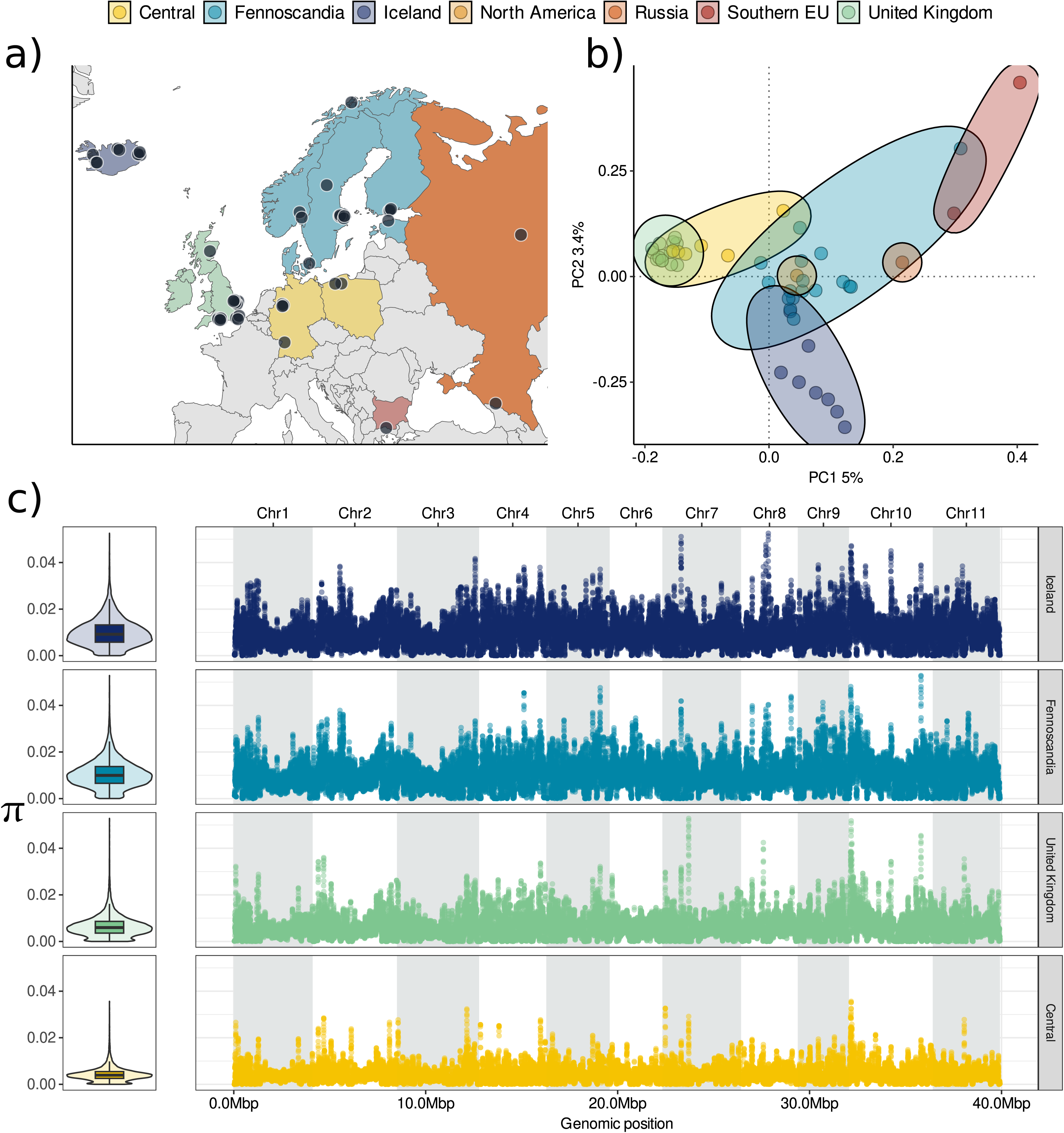
Population structure and genetic diversity of the European lineage of *B. edulis*. a) Sampling locations of 49 genets, colour-coded by geographical region. b) Scatter plots of individual variation in principal component (PC) scores derived from principal component analysis (PCA) of the genomic data. The amounts of variation explained by each PC are given as percentages on the axis labels, and the samples are color coded as shown in panel a). c) Nucleotide diversity (π) calculated in 10kb sliding windows across the genome for every sub-population. The left panels show violin plots of the kernel densities of π together with standard Tukey boxplots (centre line = median, bounds of the box = 25^th^ and 75% percentiles, upper and lower whiskers = largest and smallest value but no further than 1.5 * inter-quartile range from the hinge). The right panels show π values for each window across the 11 chromosomes. The dashed white lines represent lineage-specific genome-wide average π values.

Genetic diversity was not evenly distributed among the genetic clusters (Figure 4b). Populations at higher latitudes, specifically those in Iceland and Fennoscandia, exhibited higher genome-wide nucleotide diversity (π) than populations from the United Kingdom and Central Europe. This observation aligns with our previous finding of a negative (but non-significant) association between latitude and genomic inbreeding within the European *B. edulis* lineage (Brejon Lamartinière et al. 2024). Several factors may contribute to this pattern, including reduced habitat fragmentation due to human infrastructure in northern populations, where much of Iceland and Fennoscandia remains covered by forests or tundra. Additionally, differences in the amount of gene flow with the neighboring ‘AK’ lineage, which spans Alaska and Siberia, could also play a role. Above and beyond this pattern, we also identified regions of elevated nucleotide diversity within the *B. edulis* genome. Some of these regions were shared among the clusters, such as a prominent peak near the beginning of chromosome ten, while others were not universally shared, appearing only on specific chromosomes (Figure 4b). These regions could represent areas of structural variation, they may be under balancing selection, or they might experience locally elevated recombination. This highlights the potential of the European lineage of *B. edulis* as a model for studying these diverse evolutionary processes.

## Conclusion

Population genomic studies of EMF hold great potential for advancing our understanding of these essential organisms. However, the contiguity of reference genomes continues to be a limiting factor. This study presents one of the first haplotype-resolved chromosomal reference assembly for a dikaryotic EMF, achieving near telomere-to-telomere coverage across multiple chromosomes. Using chromatin conformation capture, we successfully anchored PacBio long reads to chromosomes for each haplotype. This approach revealed structural variation within homologous chromosomes of the reference individual and identified key areas for future research. Specifically, we identified mating loci, CAZymes and effector proteins, discovered a 0.4 Mb inversion on chromosome 3, investigated the population genetic structure of the European lineage, and discovered several genomic regions with locally elevated nucleotide diversity. We anticipate that this new resource will facilitate future discoveries in this enigmatic and ecologically important fungus.

## Data availability

The haplotype assemblies and short reads are available at NCBI under the bioproject PRJNA1187522. The annotation files and code are deposited at zenodo (https://doi.org/10.5281/zenodo.14311982) and github https://github.com/ebrejonl/Chromosomal_reference_Boletus_edulis_EU/.

## Acknowledgments

We are grateful to BMK genomics for providing details of their methods. We thank Kosmas Hench for providing advice on coding and reproducibility, and Jinhua Zhang for sharing with us the *Suillus bovinus* genome.

## Author contributions

E.B.L. & J.I.H. conceived the study. J.I.H. acquired funding and supervised the PhD student (E.B.L.) with additional scientific input from K.K.D. and B.T.M.D. E.B.L. and K.T. analyzed the data. E.B.L. and J.I.H. drafted the manuscript. All of the authors commented upon and approved the final manuscript.

